# Mass spectrometric profiling of HLA-B44 peptidomes provides evidence for tapasin-mediated tryptophan editing

**DOI:** 10.1101/2023.02.26.530125

**Authors:** Amanpreet Kaur, Avrokin Surnilla, Anita J. Zaitouna, Venkatesha Basrur, Michael B. Mumphrey, Irina Grigorova, Marcin Cieslik, Mary Carrington, Alexey I. Nesvizhskii, Malini Raghavan

## Abstract

Activation of CD8^+^ T cells against pathogens and cancers involves the recognition of antigenic peptides bound to human leukocyte antigen (HLA) class-I proteins. Peptide binding to HLA class I proteins is coordinated by a multi-protein complex called the peptide loading complex (PLC). Tapasin, a key PLC component, facilitates the binding and optimization of HLA class I peptides. However, different HLA class I allotypes have variable requirements for tapasin for their assembly and surface expression. HLA-B*44:02 and HLA-B*44:05, which differ only at residue 116 of their heavy chain sequences, fall at opposite ends of the tapasin-dependency spectrum. HLA-B*44:02 (D116) is highly tapasin-dependent, whereas HLA-B*44:05 (Y116) is highly tapasin-independent. Mass spectrometric comparisons of HLA-B*4405 and HLA-B*44:02 peptidomes were undertaken to better understand the influences of tapasin upon HLA-B44 peptidome compositions. Analyses of the HLA-B*44:05 peptidomes in the presence and absence of tapasin reveal that peptides with the C-terminal tryptophan residues and those with higher predicted binding affinities are selected in the presence of tapasin. Additionally, when tapasin is present, C-terminal tryptophans are also more highly represented among peptides unique to B*44:02 and those shared between B*44:02 and B*44:05, compared with peptides unique to B*44:05. Overall, our findings demonstrate that tapasin influences the C-terminal composition of HLA class I-bound peptides and favors the binding of higher affinity peptides. For the HLA-B44 family, the presence of tapasin or high tapasin-dependence of an allotype results in better binding of peptides with C-terminal tryptophans, consistent with a role for tapasin in stabilizing an open conformation to accommodate bulky C-terminal residues.

## Introduction

Human Leukocyte Antigen Class I (HLA class I) molecules help the immune system identify infected and transformed cells to initiate immune responses via CD8^+^ T cells and natural killer cells (1). HLA class I molecules have peptide binding sites onto which peptide fragments of proteins are bound (2). HLA class I-peptide complexes are then presented on the cell surface, where they can trigger CD8^+^ T cell cytokine production and cytotoxicity, following specific binding to T cell receptors. Thousands of HLA class I polymorphisms exist in the human population (3), with each variant containing a unique peptide binding site that determines its peptide repertoire specificity (4). Most humans express 6 different HLA class I molecules on the surface of all nucleated cells. The peptide binding specificities of individual HLA class I variants determine their abilities to induce CD8^+^ T cell responses and contribute to adaptive immunity against viruses and cancers.

Following the processing of cytosolic proteins via the proteasome, the assembly of HLA class I molecules with peptides occurs in the endoplasmic reticulum (ER) lumen with the aid of the peptide-loading complex (PLC) (5, 6). The translocation of peptides from the cytosol into the ER lumen is mediated by the transporter associated with antigen processing (TAP), a PLC component. Tapasin, another component of the PLC, has multiple roles in HLA class I assembly. Together with the chaperone calreticulin and the oxidoreductase ERp57, tapasin interacts with peptide-deficient versions of MHC class I molecules in the ER lumen and stabilizes this conformation (6–12). In doing so, the PLC increases the peptide binding to MHC class I molecules. Tapasin also optimizes the MHC class I peptide repertoire towards the binding of higher-affinity peptides (7, 9, 10).

HLA class I allotypes are known to vary in their requirements for tapasin, exhibiting a wide spectrum of tapasin-dependencies (13–16). Noteworthy for such differences are two closely related HLA-B allotypes-HLA-B*44:05 and HLA-B*44:02, which differ by a single amino acid at position 116 within their heavy chain sequences. B*44:02 and B*44:05 are at the extreme ends of the tapasin-dependency spectrum (7, 14–16). These two allotypes could allow for a deeper understanding of the influences of tapasin on HLA class I peptidome compositions. Recent advances in mass spectrometry of HLA class I peptidomes from monoallelic cell lines have allowed for the generation of large peptidome datasets for a large number of individual HLA class I allotypes (17, 18). We applied a similar approach to examine the key differences between HLA-B*44:05 peptidomes isolated from B*44:05 monoallelic cells expressing or lacking tapasin. The marked reduction in HLA-B*44:02 expression in cells lacking tapasin precluded an analysis of the B*44:02 peptidome from cells lacking tapasin. However, the peptidomes of HLA-B*44:05 and B*44:02 from tapasin-sufficient cells were compared, revealing key insights into the effects of tapasin dependence of an allotype on peptide selection.

## Results

### Expression and purification of HLA-B*44:05 from wild type (WT) and tapasin knockout (TPN-KO) 721.221 cells

As noted above, HLA-B*44:05 is relatively tapasin-independent and detectable at significant levels on the surface of cells lacking tapasin (7, 14–16). This allotype was thus chosen for comparisons of HLA class I peptidome characteristics in the absence and presence of tapasin. The contribution of tapasin in shaping HLA class I peptidomes was determined by using CRISPR/Cas9-based gene editing to knockdown the expression of tapasin in HLA class I-deficient 721.221 cells that had been transduced to generate monoallelic 721.221-HLA-B*44:05 cells. Single-cell cloning using the limiting dilution method resulted in the generation of 721.221-B*44:05 clones with a complete tapasin knockout (B*44:05-tapasin-KO) (**Figure 1A**). Quantitative flow cytometry assays following surface staining by the peptide-HLA class I complex-specific W6/32 antibody (19, 20) showed a significant reduction in the number of HLA-B*44:05 molecules on the surface of 721.221-B*44:05-tapasin-KO (B*44:05-TPN-KO) cells when compared to wild type 721.221-B*44:05 (B*44:05-WT) cells (**Figure 1B**). In order to determine if reduced stability could account for lower cell surface B*44:05 in B*44:05-tapasin-KO cells, we analyzed surface B*44:05 expression on B*44:05-WT *vs*. B*44:05-tapasin-KO cells after blocking the trafficking of newly synthesized HLA-B molecules for various time (up to 4h) using Brefeldin A. We observed a trend towards a decrease in the stability of surface HLA-B*44:05 expression upon the knockdown of tapasin (**Figures 1C and 1D**). The calculated average half-life of surface B*44:05 is reduced from ~6.22 hours in B*44:05-WT cells to ~4.7 hours in B*44:05-tapasin-KO cells. There were rather large variations in the half-life values of HLA-B*44:05 on the surface of B*44:05-WT cells, based on multiple replicates (ranging from ~3.6 hours to ~10.3 hours, **Figure 1D**).

**Figure 1:**
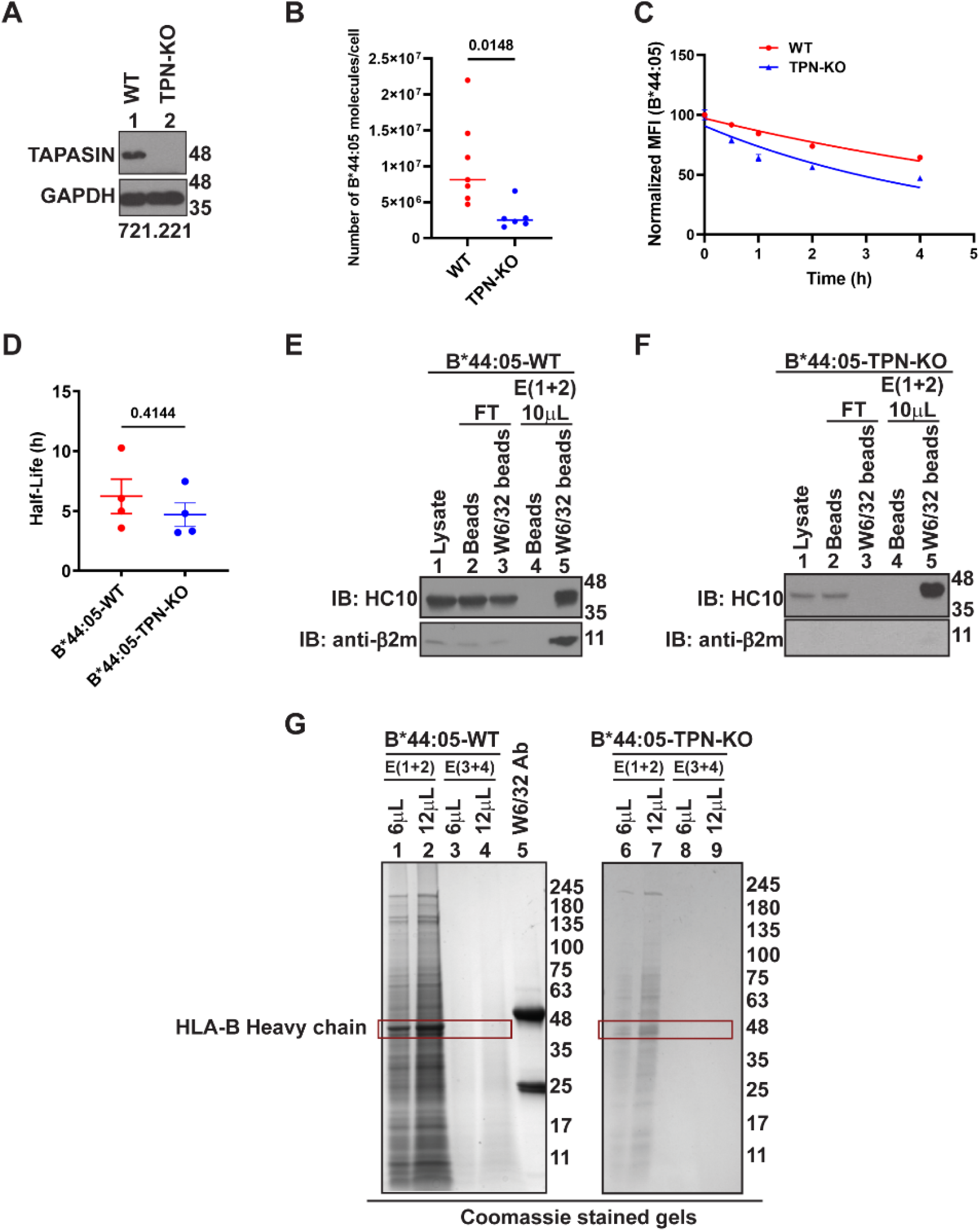
Expression and purification of HLA-B*44:05 from wild type (WT) and tapasin knockout (TPN-KO) 721.221 cells: **(A)** Representative blots showing tapasin (TPN) expression in the lysates of 721.221-B*44:05 WT (lane 1) and tapasin knockout (lane 2) cells. The sgRNA sequence is given in the materials and methods. **(B)** The graph shows the mean of the number of HLA-B*44:05 molecules expressed per cell on the surface of WT (n=7) and TPN-KO (n=6) 721.221-B*44:05 cells measured by quantitative flow cytometry following staining with W6/32-FITC antibody. The number of HLA-B molecules per cell is quantified from standard curves of the geometric mean fluorescence intensity (MFI) values of W6/32-FITC antibody binding to Quantum Simply Cellular beads (Bangs Laboratory, Inc.) that contain a known number of Fc receptors and are stained concurrently with the cells. Each data point represents an independent experiment. Statistical significance was calculated using an unpaired t-test assuming equal variance. **(C)** Representative plot shows changes in the expression (normalized relative to the 0h condition) of B*44:05 on the surface of WT and TPN-KO 721.221-B*44:05 cells, assessed using the W6/32 antibody, at different time points following brefeldin A treatment. **(D)** The graph shows calculated half-lives of B*44:05 on the surface of B*44:05-WT and B*44:05-TPN-KO 721.221 cells based on the surface stability measurements shown in C (n=4). Statistical significance is calculated using a two-tailed unpaired t-test. Graphs are plotted in GraphPad Prism. **(E and F)** Representative blots (of six independent experiments) show the purification of HLA-B*44:05 from B*44:05-WT (E) and B*44:05-TPN-KO 721.221 cells (F). “Beads” indicates protein A beads used for pre-clearing of lysates and “W6/32 beads” indicates protein A beads crosslinked to W6/32 antibody. HLA-B heavy chain and β2m are probed using HC10 and anti-β2m antibodies, respectively. **(G**) Representative Coomassie brilliant blue stained gels (of six independent experiments) showing elution of B*44:05 from W6/32 beads following affinity purification from B*44:05-WT or B*44:05-TPN-KO 721.221 cells. The red outlined box marks the HLA-B heavy chain bands in the eluates. E, Elution; FT, Flow through; IB, Immunoblot

For the peptidome analyses, we adopted an immunoaffinity (IA)-based approach for the purification of peptide-HLA-B complexes followed by LC-MS/MS-based identification of HLA-B binding peptides (17, 18, 21). The W6/32 antibody was covalently crosslinked to Protein A beads and used for the purification of peptide-HLA class I complexes from B*44:05-WT or B*44:05-tapasin-KO 721.221 cells. The successful purification of peptide-HLA class I complexes from the lysates of 721.221 cells was confirmed via western blots using anti-heavy chain (HC10) (22) and anti-β2m antibodies (**Figures 1E and F**). The presence of heavy chain and β2m signals in the elutions from W6/32 beads (**Figure 1E and Figure 1F; lane 5**) in contrast to the elutions from blank beads (**Figure 1E and Figure 1F; lane 4**) further confirmed the successful purification of HLA class I. Relative to the heavy chain band intensity, the β2m signal was low for B*44:05 purified from B*44:05-tapasin-KO cells compared to that from B*44:05-WT cells (**Figures 1E and F, lane 5, top and lower panels**), again indicating the reduced stability of HLA-B*44:05 complexes in the absence of tapasin. In the Coomassie blue stained SDS-PAGE gels, we observed a strong band at a size consistent with that of HLA class I heavy chains (**~48 kDa**) in the elutions from W6/32 beads, which migrated at a distinct size from the antibody heavy chain (**Figure 1G, lanes 1 and 2 compared with lane 5**). Additionally, the overall amount of HLA class I heavy chain recovered was reduced in the elution fractions from W6/32 beads following B*44:05 purification from B*44:05-tapasin-KO cells compared to that from B*44:05-WT cells **(Figure 1G).** The HLA class I bound peptides were further fractionated from the purified proteins using C18 reversed-phase chromatography and used for mass spectrometric analyses.

### Greater non-canonical characteristics of B*44:05 peptides from tapasin-KO cells

Peptides identified from LC-MS/MS analysis of elutions from W6/32 beads were curated by selection for peptide lengths of 8 to 14 amino acids. Non-specific peptides were removed as described in the methods. The number of peptides identified in six individual LC-MS/MS runs ranged from 701 to 3259 (median value of 2187) for B*44:05-WT cells vs. 1093 to 3245 (median value of 1533) for B*44:05-tapasin-KO cells. For immunopeptidome characterizations, peptides identified in the six independent LC-MS/MS runs of B*44:05 eluates from WT and tapasin-KO cells were compared. The peptides that were overlapping across ≥ 2 runs were chosen for further comparisons of their characteristics. A total of 1559 and 2066 peptides were identified for the B*44:05-WT and B*44:05-tapasin-KO conditions, respectively (**Figure 2A**). Of these, 272 peptides were unique to the B*44:05-WT, 779 peptides were unique to the B*44:05-tapasin-KO condition, while 1287 peptides were shared between the two conditions (**Figure 2A**). The tapasin-KO group has a larger pool of unique peptides, suggesting that this condition is tolerant of more diverse sequences.

**Figure 2:**
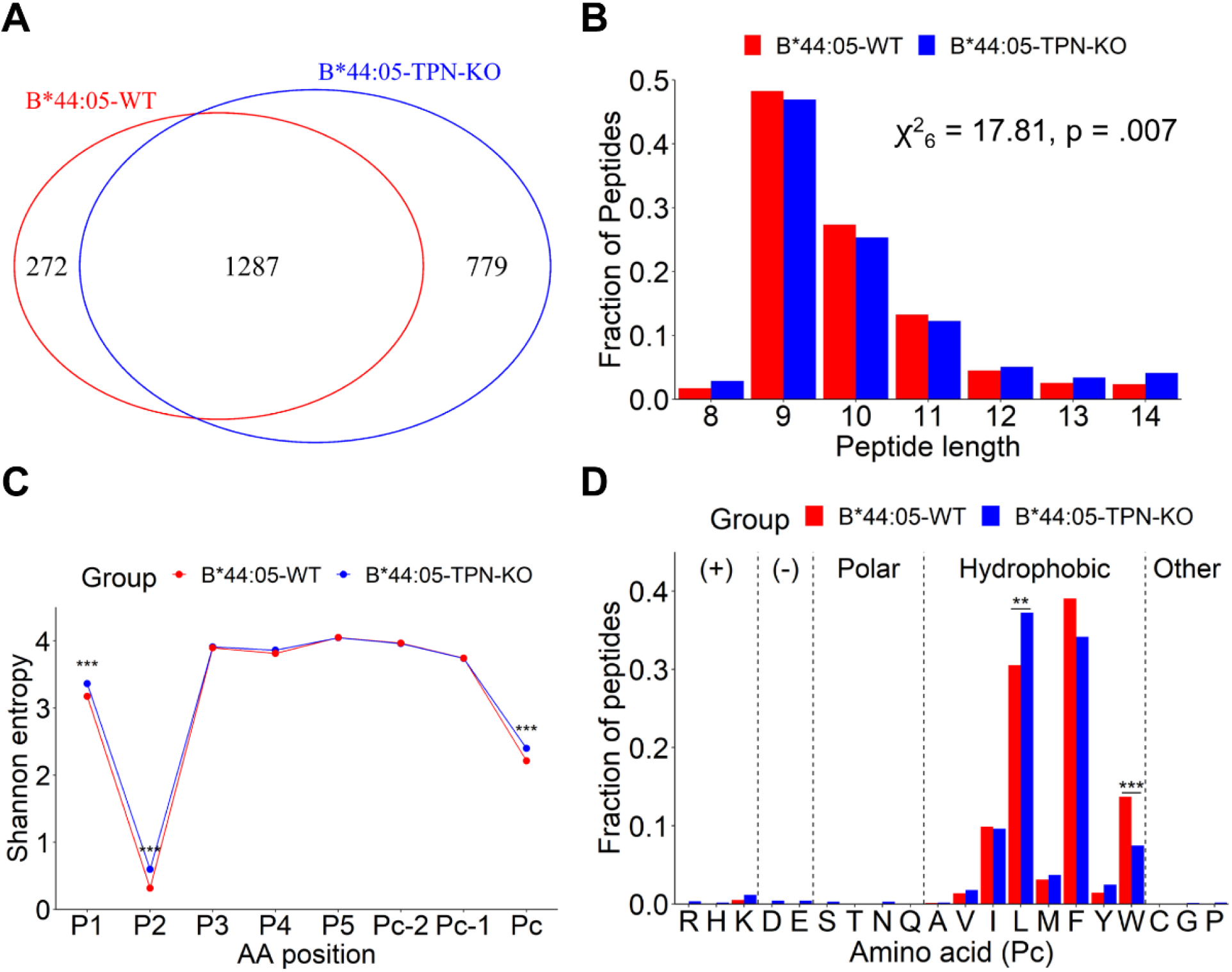
Charateristics of HLA-B*44:05 peptides purified from wild type or tapasin-KO cells. **(A)** Venn diagram showing total number of peptides identified from six MS/MS runs grouped into the unique to B*44:05-TPN-KO, unique to B*44:05-WT, or shared categories. Only peptides observed in 2 or more runs are included. **(B)** Length distributions of peptides identified for the B*44:05-WT and B*44:05-TPN-KO categories, indicated as a fraction of total number of peptides. Significant differences in distribution were determined using the Chi-squared test of independence. **(C)** Shannon Entropy (SE) plots for indicated positions within the 9-mer, 10-mer, and 11-mer peptides from (A). The P_C_, P_C-1_ and P_C-2_ represent the C-terminal, and the −1 and −2 positions relative to the C-terminus, respectively. Significant differences in SE at each position were determined using a bootstrap hypothesis test with Bonferroni corrected p-values. **(D)** The fractional distribution of all amino acids at the P_C_ position of 9-mer, 10-mer, and 11-mer peptides, grouped according to the side chain properties of individual amino acids. Significant differences in the amino acid distributions were determined using the Chi-squared test of independence, with standardized residual post-hoc tests using Benjamini-Hochberg corrections. *: p < 0.05; **: p < .01; ***: p < .0001

The most frequent peptide lengths observed were 9-11-mers for both B*44:05-WT or B*44:05-tapasin-KO cells, as expected for HLA class I-binding peptides (**Figures 2B**). However, 9-11-mers constituted a smaller fraction of B*44:05 peptides under the tapasin-KO condition when compared to the wild type condition (**Figures 2B**). Non-canonical lengths were more prevalent in the tapasin-KO condition and the overall length distribution differences were statistically significant (**Figure 2B**; *χ*^2^ =17.81; p value=0.007).

Peptide diversity analysis using Shannon entropy (SE) calculations (23, 24) indicated the lowest entropy and strongest restriction at the P_2_ position for both the B*44:05-WT and B*44:05-tapasin-KO peptide groups (**Figure 2C**). The P_C_ position was the second most restrictive position across different peptide lengths for all the groups (**Figures 2C**). The strong restrictions observed at the P_2_ and P_C_ positions are consistent with the previous reports that have described P_2_ and P_C_ amino acids as anchor residues for binding of peptide ligands to the members of the HLA-B44 supertype (25). Comparative analyses of SE plots of 9-11-mers suggest that the peptides identified under B*44:05-tapasin-KO condition are significantly less restrictive at the P_1_, P_2_ and P_C_ positions compared to the peptides identified under B*44:05-WT condition (**Figure 2C**). Overall, these data suggest that tapasin deficiency reduces the restrictiveness of peptide binding specificities at the main anchor positions within the peptide binding groove of the B*44:05 allotype.

### Reduced tryptophan prevalence in the P_C_ position of B*44:05 peptides purified from tapasin-KO cells

The fractional distribution of all twenty amino acids at P_C_ position of 9-11-mer B*44:05 peptides in both the wild type and tapasin-KO groups revealed a strong preference for hydrophobic amino acids with a high prevalence of leucine and phenylalanine followed by isoleucine and tryptophan (**Figure 2D**). Comparative analysis revealed a significant increase in the fraction of peptides with leucine at the P_C_ position among the 9-11-mers in the B*44:05-tapasin-KO group when compared to 9-11-mers in the B*44:05-WT group (**Figure 2D**). On the other hand, there was a significant decrease in the representation of peptides with tryptophan at the P_C_ position among 9-11-mer B*44:05 peptides identified under the tapasin-KO condition compared to those in the B*44:05-WT group (**Figure 2D**).

The identified peptides were further categorized into unique and shared groups, based on their presence only in one of the WT or the tapasin-KO conditions (unique peptides, not observed in any run of the other condition) or shared across both conditions (shared peptides) (**Supplementary table 1**). Seq2Logo plots (26) were generated to further examine the residue preferences of B*44:05 and to better understand the influences of tapasin on the peptidome compositions. The Seq2Logo motifs for 9-11-mer peptides that are unique to either B*44:05-WT or B*44:05-tapasin-KO conditions or shared across the two conditions revealed a strong preference for glutamic acid at the P_2_ position (**Figure 3**). Consistent with the results of fractional distribution of amino acids at P_C_ position (Figure 2D), the peptides unique to the B*44:05-WT condition (**Figure 3A**) displayed a higher tryptophan prevalence at the P_C_ position relative to the peptides unique to the B*44:05-tapasin-KO condition (**Figure 3B**) or the shared peptides (**Figure 3C**). While the C-terminal tryptophans were present in the shared peptides group (**Figure 3C**), as well as in the total B*44:05 peptides (including the shared and unique peptides) from the tapasin-KO cells (**Figure 3E**), the preference for C-terminal tryptophan was more pronounced among the peptides isolated from B*44:05-WT cells (unique (**Figure 3A**) and total (**Figure 3D**)).

**Figure 3:**
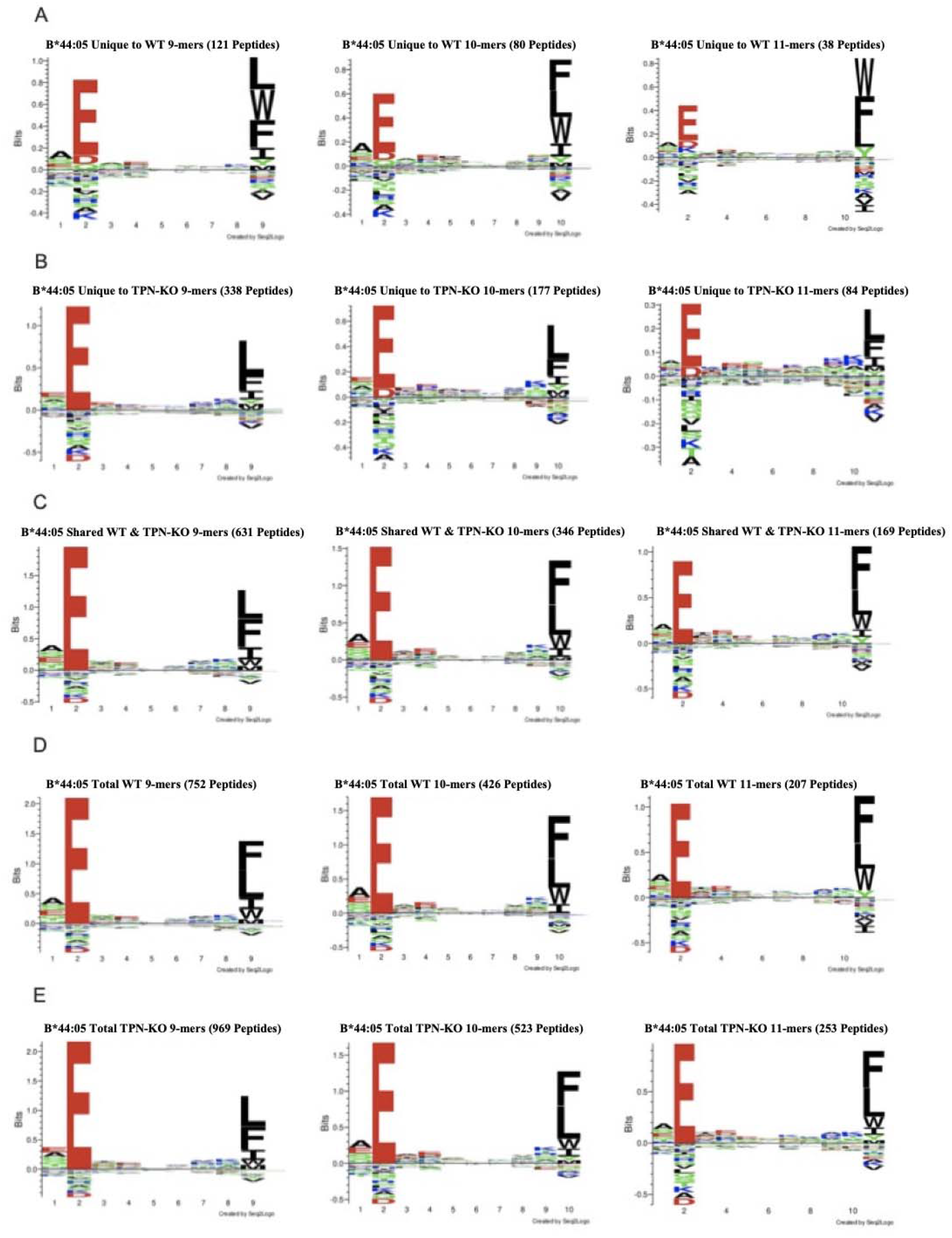
Peptides unique to B*44:05 from 721.221-WT cells show increased preference for tryptophan at the C-terminus: **(A to C)** Seq2Logo motifs for 9-mer, 10-mer, and 11-mer peptides from the peptide set highlighted in Figure 2A, grouped into the unique to B*44:05-WT (A), unique to B*44:05-TPN-KO (B) and peptides shared across the two conditions (C). **(D and E)** The Seq2Logo motifs for the total peptides (including the unique and shared peptides) for the B*44:05-WT (D) and B*44:05-TPN-KO (E) conditions are also shown.

### Peptides unique to tapasin-KO condition were predicted as weak B*44:05 binders

Next, we predicted the binding affinities of peptides for B*44:05 using NETMHCpan 4.1 (27) for 9-mer, 10-mer, and 11-mer peptides observed in the unique and shared groups. Interestingly, 11-mer peptides, across all the peptide groups, had the lowest predicted binding affinities (**Figure 4,** a higher number on Y-axis corresponds to a lower predicted binding affinity). Across all three peptide lengths, the peptides unique to the B*44:05-tapasin-KO condition yielded the lower predicted binding affinities (**Figure 4**). On the other hand, the peptides unique to the B*44:05-WT condition were predicted to have higher binding affinities across all three peptide lengths. The shared peptides were predicted to have intermediate affinities relative to the two unique groups (**Figure 4**). Lower predicted binding affinities of peptides unique to the B*44:05-tapasin-KO condition further substantiate the reduced restrictiveness of peptide binding to B*44:05 receptors in the tapasin-KO cells.

**Figure 4.**
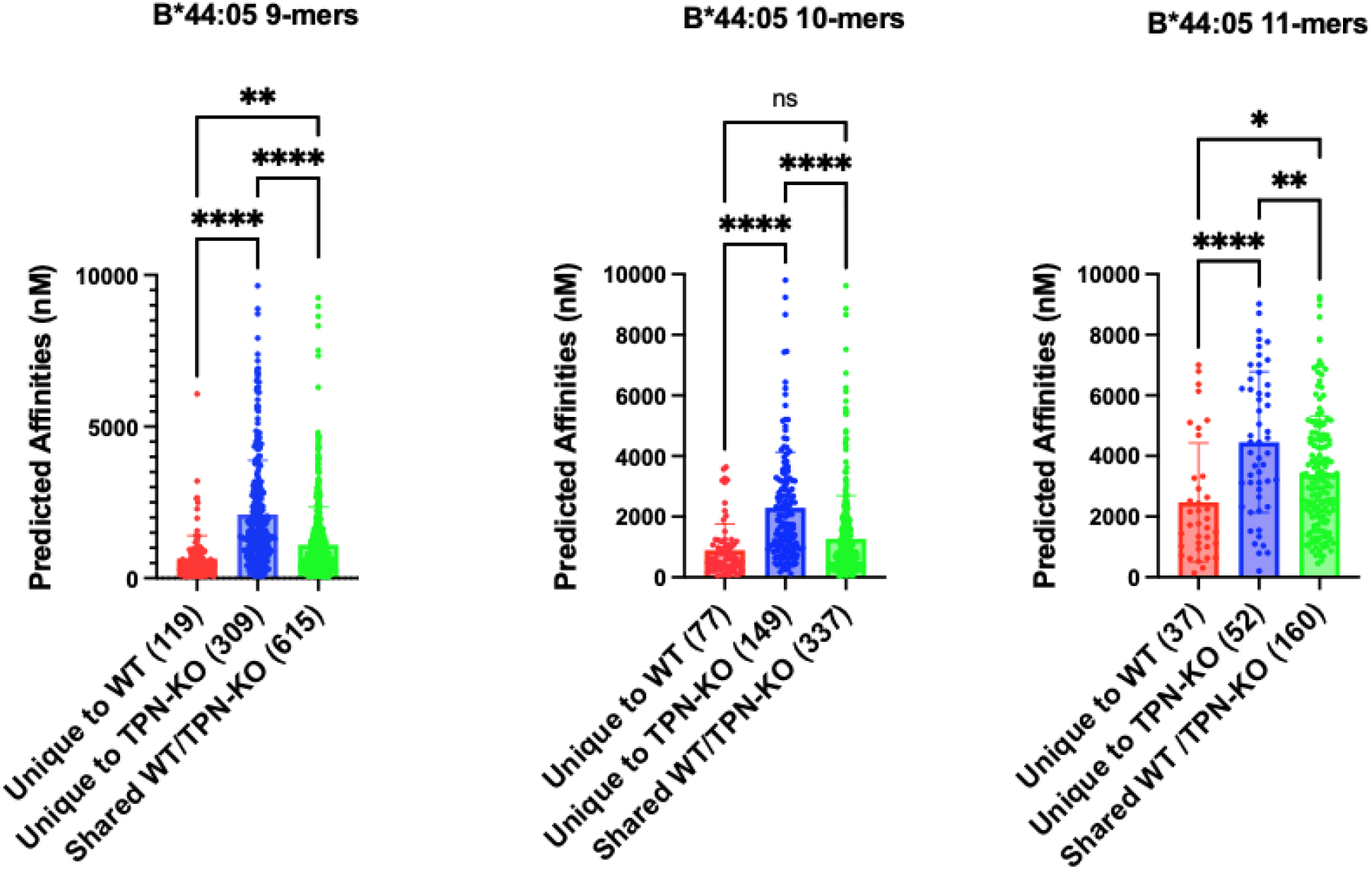
Tapasin knockout results in reduced predicted binding affinities of peptides bound to B*44:05: The predicted binding affinities (nM) of B*44:05 peptides were calculated for the indicated peptide groups and plotted separately for the 9-mer, 10-mer, and 11-mer peptides. Peptides with predicted affinities >10,000 nM were excluded.

### Expression and purification of HLA-B*44:02 from 721.221 cells, and comparison of surface expression and stability of HLA-B*44:02 and HLA-B*44:05

Next, we compared the peptidomes of B*44:02 and B*44:05 allotypes, which as previously stated, vary markedly in their tapasin dependencies (7, 14–16) despite a single difference at residue 116 in their F-pockets. Quantitative flow cytometry of surface staining with W6/32 antibody showed, on average, a lower expression of HLA-B*44:02 on the surface of 721.221 WT cells when compared to HLA-B*44:05 (p=0.013, **Figure 5A**). Despite higher surface expression, B*44:05 had a trend towards lower cell surface stability than B*44:02 (p=0.072, **Figure 5B**). Based on the measurements of the surface stabilities, the average half-life of B*44:02 surface expression (9.72 hours) was slightly higher than the average half-life of B*44:05 surface expression (7.04 hours) (**Figure 5C**).

**Figure 5:**
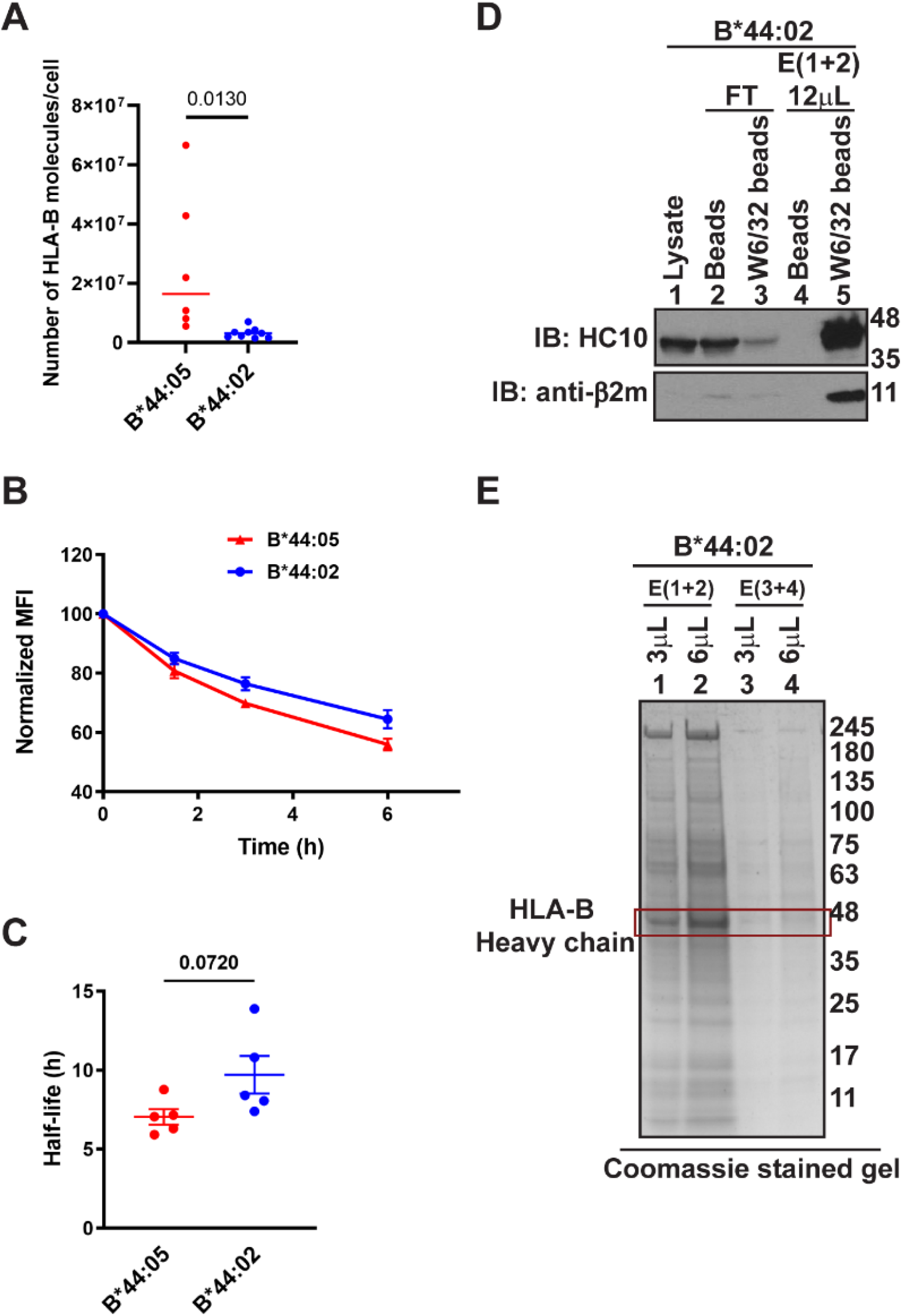
Expression and purification of HLA-B*44:02 from 721.221 cells: **(A)** The graph shows the mean number of HLA-B molecules expressed per cell on the surface of B*44:02 (n=9 independent experiments) and B*44:05 (n=6 independent experiments) 721.221 cells measured by quantitative flow cytometry following staining with W6/32-FITC antibody. The number of HLA-B molecules per cell is quantified from standard curves of the geometric mean fluorescence intensity (MFI) values of W6/32-FITC antibody binding to Quantum Simply Cellular beads (Bangs Laboratory, Inc.) that contain a known number of Fc receptors and are stained concurrently with the cells. Each data point represents an independent experiment. Statistical significance was calculated using an unpaired t-test assuming equal variance. **(B)** Representative plot shows changes in the expression (normalized relative to the 0h condition) of HLA-B on the surface of 721.221-B*44:02 and 721.221-B*44:05 cells assessed using the W6/32 antibody, at different time points following brefeldin A treatment of cells. **(C)** The graph shows calculated half-lives of B*44:02 and B*44:05 on the surface of 721.221 cells based on the surface stability measurements shown in B (n=4). Statistical significance is calculated using a two-tailed unpaired t-test. Graphs are plotted in GraphPad Prism. **(D)** Representative blots (of six independent experiments) show the purification of HLA-B*44:02 from 721.221 cells. “Beads” indicates protein A beads used for pre-clearing of lysates and “W6/32 beads” indicates protein A beads crosslinked to W6/32 antibody. HLA-B heavy chain and β2m are probed using HC10 and anti-β2m antibodies, respectively. **(E)** Representative Coomassie brilliant blue stained gel (of six independent experiments) showing elution of HLA-B*44:02 from W6/32 beads following affinity purification from 721.221-B*44:02 cells. The red outlined box marks the HLA-B heavy chain bands in the eluates. E, Elution; FT, Flow through; IB, Immunoblot

Surface peptide-MHC-I complexes were purified from 721.221-B*44:02 cells using Protein A beads conjugated to W6/32 antibody in the same manner as explained for B*44:05 (Figure 1). The presence of HC10 and β2m signals in the immunoblots of elutions from W6/32 beads (**Figure 5D; lane 5**) in contrast to the elutions from blank beads **(Figure 5D; lane 4)** confirmed the purification of HLA-B*44:02. A band at the specific size of HLA-B*44:02 heavy chain (**~48 kDa; Figure 5E**) was observed in the eluates of the W6/32 beads in the Coomassie blue stained SDS-PAGE gels. The HLA-B*44:02 bound peptides were further fractionated from the purified proteins using C18 reversed-phase chromatography and used for mass spectrometric analyses.

### Distinct restrictiveness and C-terminal preferences among peptides bound to B*44:02 and B*44:05

Peptides that were identified by LC-MS/MS analysis following purification of peptide-MHC-I complexes from B*44:02 and B*44:05 cells were first curated for length and specificity as described in Figure 2. The number of peptides identified in the six individual LC-MS/MS runs ranged from 635 to 3187 (median value of 2413) for the B*44:05 condition and from 711 to 3085 (median value of 1065) for the B*44:02 condition. Peptides identified in the six-independent LC-MS/MS runs of eluates from either B*44:05 or B*44:02 cells were included for the immunopeptidome characterizations and comparisons. The peptides that were overlapping across ≥ 2 runs were chosen for subsequent comparative analyses. A total of 1722 and 1526 peptides were respectively identified for the B*44:05 and B*44:02 conditions (**Figure 6A**). Of these, 1049 peptides were unique to the B*44:05, 853 peptides were unique to the B*44:02, while 673 peptides were shared between the two conditions (**Figure 6A**). The most common peptide lengths observed among the peptides identified for both B*44:05 and B*44:02 conditions were 9-11-mers. There were no significant differences in the length distributions of peptides identified for B*44:05 and B*44:02 conditions (**Figure 6B**; *χ*^2^ =6.06; p value=0.42).

**Figure 6:**
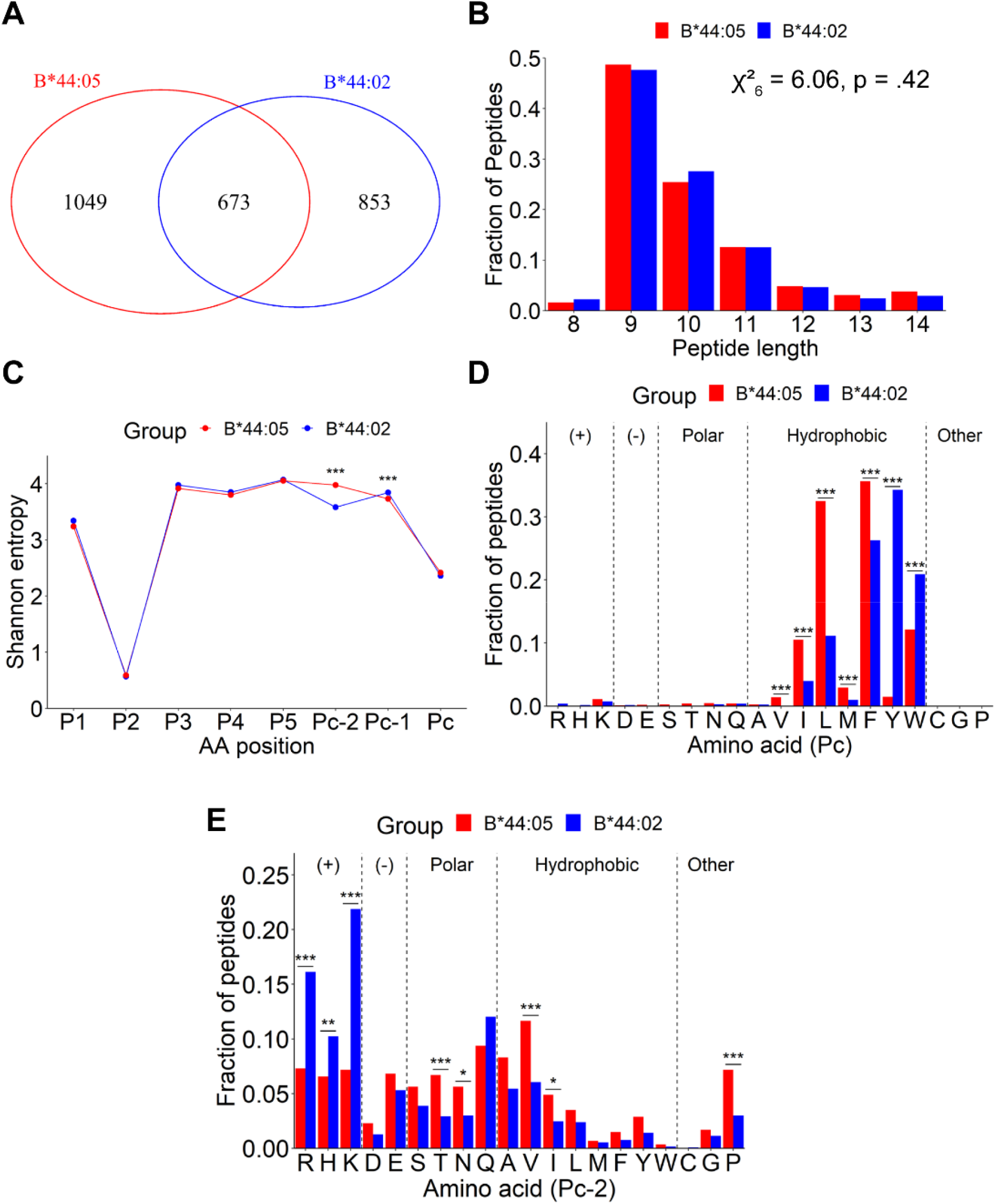
Characteristics of peptides purified from B*44:02 compared with those from B*44:05. **(A)** Venn diagram showing the total number of peptides identified from six MS/MS runs grouped into those unique to B*44:02, unique to B*44:05, or shared between the two categories. Only peptides observed in 2 or more runs are included for further analyses. **(B)** Length distributions of identified peptides, indicated as a fraction of total number of peptides. Significant differences in distribution were determined using the Chi-squared test of independence. **(C)** Shannon Entropy (SE) plots for indicated positions within 9-mer, 10-mer, and 11-mer peptides from (A). The P_C_, P_C-1_ and P_C-2_ represent the C-terminal, and the −1 and −2 positions relative to the C-terminus, respectively. Significant differences in SE at each position were determined using a bootstrap hypothesis test with Bonferroni corrected p-values. **(D, E)** The fractional distribution of all amino acids at the P_C_ (D) and P_C-2_ (E) positions of 9-mer, 10-mer, and 11-mer peptides. Peptides are grouped by side chain properties. Significant differences in distributions were determined using the Chi-squared test of independence, with standardized residual post-hoc tests using Benjamini-Hochberg corrections. *: p < 0.05; **: p < .01; ***: p < .0001

Peptidome diversity analysis using SE calculations indicated the lowest entropy and strongest restriction at the P_2_ position for both B*44:05 and B*44:02 peptides. The P_C_ position was the second most restricted position across the two peptide groups (**Figure 6C**). Comparative analyses of the SE plots of 9-11-mers among B*44:02 and B*44:05 peptides showed no significant differences in the restrictiveness of peptide binding to B*44:02 versus B*44:05 at either P_2_ or P_C_ positions. Interestingly, a significantly higher restriction was seen at the P_C-2_ position for B*44:02 peptides (**Figure 6C**) which was not observed for B*44:05 peptides. On the other hand, compared to B*44:02 peptides, a small but significant decrease in SE value for P_C-1_ position was also observed among B*44:05 peptides (**Figure 6C**).

Analysis of the fractional distribution of all twenty amino acids revealed a strong preference for hydrophobic amino acids at P_C_ position among peptides identified for both B*44:02 and B*44:05 allotypes (**Figure 6D**). Among hydrophobic amino acids, a significantly higher prevalence of tryptophan or tyrosine at P_C_ position is observed among B*44:02 peptides when compared to B*44:05 peptides (**Figure 6D**). In contrast to B*44:02 peptides, B*44:05 peptides display a substantially higher preference for less bulky hydrophobic amino acids such as phenylalanine, leucine, isoleucine, methionine, and valine at P_C_ position (**Figure 6D**). The fractional amino acid distributions at P_C-2_ position varied considerably between B*44:02 and B*44:05 peptides (**Figure 6E**). The B*44:02 peptides exhibit a significant higher abundance of positively charged amino acids-histidine, arginine and lysine at P_C-2_ position when compared to B*44:05 peptides (**Figure 6E**). In contrast to B*44:02 peptides, the B*44:05 peptides display a relatively smaller but significant enrichment of uncharged aliphatic amino acids-valine, threonine, proline, asparagine, and isoleucine at P_C-2_ position (**Figure 6E**). Thus, HLA-B*44:02 and HLA-B*44:05 allotypes display distinct preferences for amino acids at P_C_ and P_C-2_ positions of peptides that bind to the two B44 allotypes.

The identified peptides were further categorized into unique and shared groups, based on their detection in either the B*44:02 or the B*44:05 condition (unique peptides, not observed in any run of the other condition) or across both conditions (shared peptides) (**Supplementary table 2**). Analyses of the Seq2Logo plots showed that the peptides unique to B*44:02 (**Figure 7B**) as well as the total B*44:02 peptides (unique + shared with B*44:05) (**Figure 7E**) display a strong preference for tyrosine at the P_C_ position. A strong preference was also observed for tryptophan at the P_C_ position among the peptides unique to B*44:02 (**Figure 7B**) as well as the full set of B*44:02 peptides (total; **Figure 7E**). However, while the shared peptides (**Figure 7C**) and the full set of B*44:05 peptides (total; **Figure 7D**) showed some tryptophan enrichment at the P_C_ position, tryptophan at the P_C_ position was largely excluded among the peptides unique to B*44:05 (**Figure 7A**). On the contrary, among the peptides unique to B*44:05 as well as the full set of B*44:05 peptides (**Figure 7A and 7D,** respectively), there was an elevated preference for leucine and phenylalanine at the P_C_ position. Additionally, the peptides unique to B*44:02 and total B*44:02 peptides (**Figure 7B and 7E,** respectively) showed an enrichment of the positively charged amino acids, lysine and arginine, at their P_C-2_ positions. This preference aligns with the higher restrictiveness observed in SE plots at the P_C-2_ position among the peptides unique to B*44:02 (Figure 6C). In contrast, the peptides unique to B*44:05 and total B*44:05 peptides (**Figure 7A and 7D,** respectively) showed no specific amino acid restrictions or preferences at their P_C-2_ positions. Interestingly, the peptides shared between B*44:02 and B*44:05 also did not exhibit any specific preferences at their P_C-2_ positions (**Figure 7C**).

**Figure 7:**
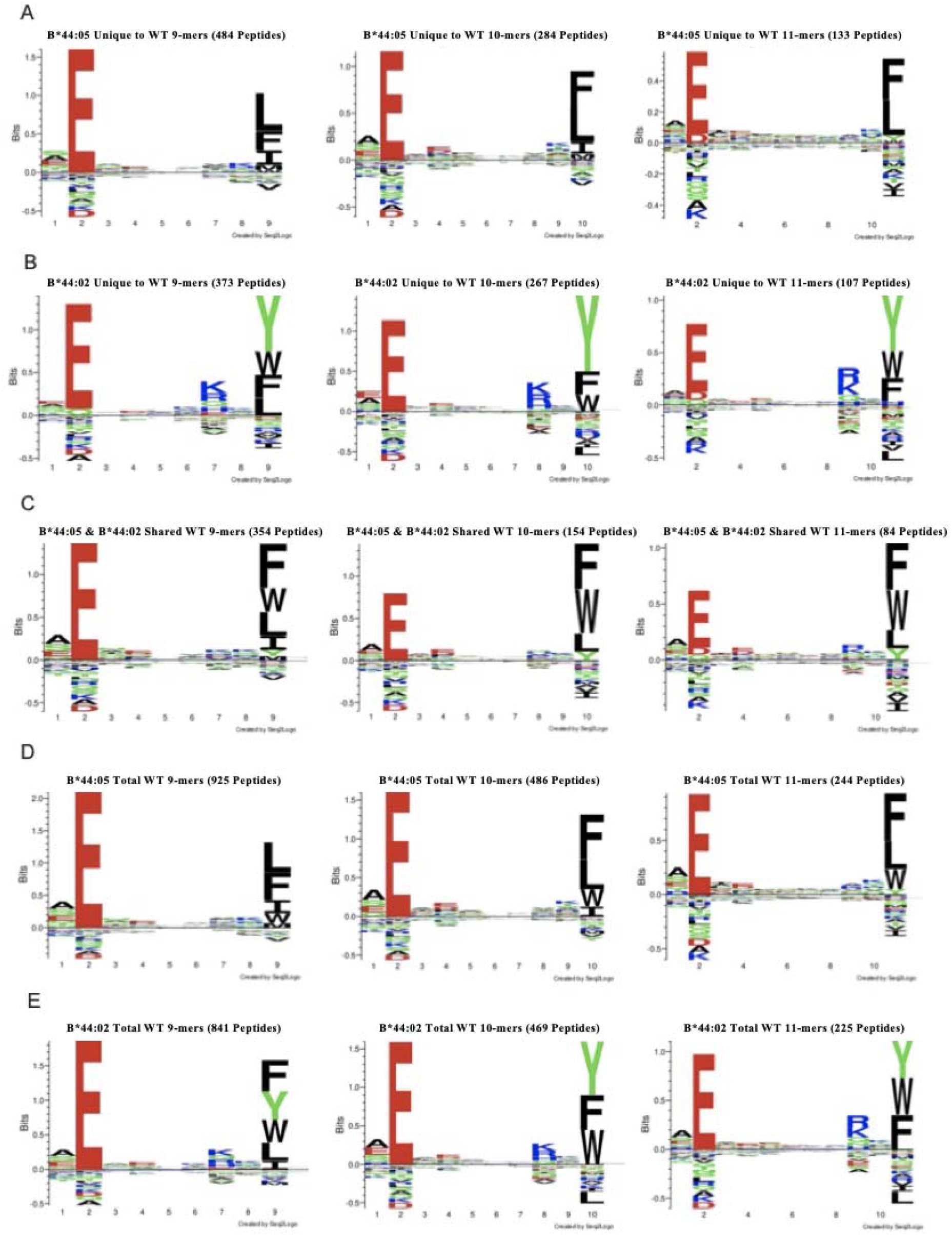
Tryptophan is under-represented among peptides unique to B*44:05 relative to those shared with B*44:02: **(A to C)** Seq2Logo motifs for 9-mer, 10-mer, and 11-mer peptides among the peptides unique to B*44:05 (A), unique to B*44:02 (**B**) and shared between B*44:02 and B*44:05 (C). **(D and E)** The Seq2Logo motifs for the total set of peptides (including the unique and shared peptides) for B*44:05 (D) and B*44:02 (E) are also shown.

We predicted the binding affinities of peptides for B*44:02 or B*44:05 using NETMHCpan 4.1 (27) for 9-mer, 10-mer, and 11-mer peptides observed in the unique and shared groups. Importantly and notably, the predicted binding affinities of B*44:05 peptides shared with B*44:02 are significantly higher than those of the peptides unique to B*44:05 (**Figure 8A;** higher numbers on Y-axis indicate lower binding affinities). On the other hand, despite the altered specificities of the peptides unique to B*44:02 compared to those shared between B*44:02 and B*44:05, no significant differences were observed in the predicted binding affinities (**Figure 8B**). These findings suggest that while the peptidomes of tapasin-independent allotypes such as HLA-B*44:05 may include both optimized and suboptimal peptides, the peptides bound to a strictly tapasin-dependent allotype such as HLA-B*44:02 are significantly optimized.

**Figure 8.**
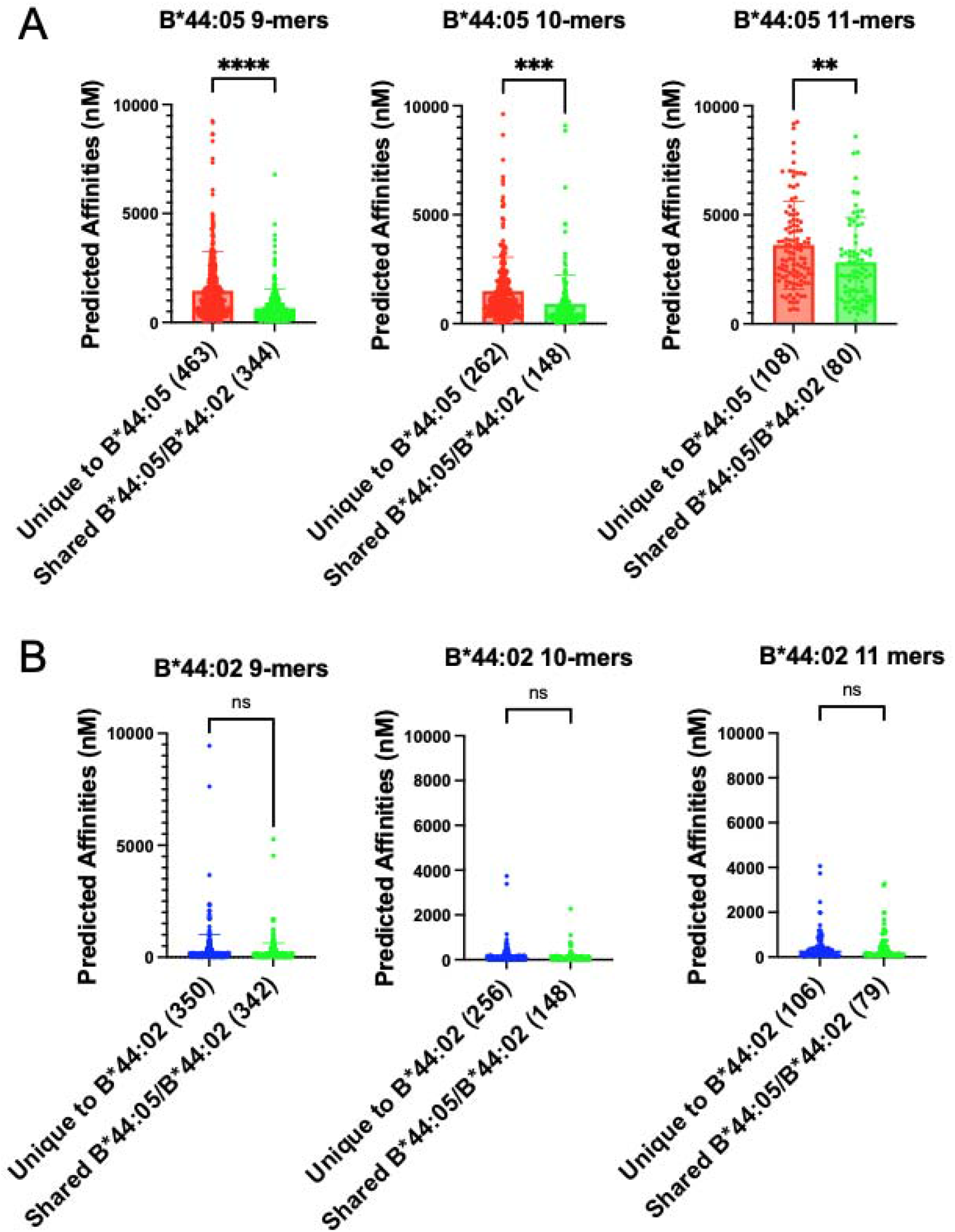
Reduced predicted binding affinities of peptides unique to B*44:05 compared to those shared with B*44:02: **(A)** The predicted affinities (nM) of B*44:05 peptides were calculated for the indicated groups, and plotted separately for the 9-mer, 10-mer, and 11-mer peptides. Peptides with predicted affinities >10,000 nM are excluded. **(B)** The predicted affinities (nM) of B*44:02 peptides were calculated for indicated groups, and plotted separately for the 9-mer, 10-mer, and 11-mer peptides. Peptides with predicted affinities >10,000 nM are excluded.

## DISCUSSION

The integral role of the PLC in antigen presentation to CD8^+^ T cells via HLA class I molecules makes the PLC a preferred target for downmodulation in different pathogenic diseases and cancers (28–30). However, the differential dependence of HLA class I allotypes on PLC components can have profound implications for the incidence and prognosis of these diseases. In this study, we compared the peptidomes of two members of the B44 supertype, HLA-B*44:02 and HLA-B*44:05, that differ significantly in their tapasin dependencies, ER retention and thermostabilities (7, 14–16, 31, 32), despite just a single amino acid difference within the heavy chain sequences. HLA-B*44:02 has a negatively charged aspartate residue at position 116 in contrast to a tyrosine residue in HLA-B*44:05.

We used CRISPR/Cas9-based gene editing to knockout the expression of tapasin in 721.221 cells expressing HLA-B*44:05. We observed a significant reduction in the surface expression of B*44:05 in tapasin-deficient cells when compared to wild-type cells (**Figure 1B**). Low surface levels of tapasin-independent B*44:05 detected under tapasin-deficient conditions could result from either the trafficking of sub-optimally assembled B*44:05 or the presence of epitope-free unstable B*44:05 heavy chains not detected by the W6/32 antibody. In favor of the former, we measured a lower stability of surface B*44:05 proteins in tapasin-deficient cells and a lower recovery of heavy chain-β2m complexes from the lysates (**Figures 1C to F**). Furthermore, the low predicted B*44:05 binding affinities for peptides grouped as unique to the tapasin-KO condition (**Figure 4**) support the model of suboptimal assembly of B*44:05 with low-affinity peptides in the absence of tapasin. Reconstitution of tapasin expression in 721.220 cells, devoid of endogenous tapasin, induces rapid maturation of HLA-B*44:05 despite no detectable incorporation of B*44:05 in PLC (14). Indeed, soluble tapasin that fails to bind TAP (33) also promotes B*44:05 maturation (14), indicating that the presence of tapasin enhances B*44:05 maturation and intracellular transport independent of PLC engagement. Tapasin-dependent optimization of B*44:05 peptide cargo has been illustrated in another study that showed an increase in the thermostability of W6/32 reactive peptide-B*44:05 complexes in the presence of tapasin (7). Our results, supported by these earlier findings, clearly demonstrate that despite the high tapasin-independence of HLA-B*44:05, the presence of tapasin facilitates optimal assembly and trafficking of more stable peptide-HLA-B*44:05 complexes to the cell surface.

The SE values for P_2_ and P_C_ positions among the peptides identified under B*44:05-tapasin-KO condition were calculated to be slightly higher than those for the B*44:05 peptides detected under the WT condition (**Figure 2C**). Importantly, we found that the peptide C-terminal tryptophan content is significantly affected by the presence of tapasin. The fraction of peptides with C-terminal tryptophan is reduced in the absence of tapasin (**Figure 2D**) and the peptides with C-terminal tryptophan are largely excluded among the peptides unique to tapasin-KO cells (**Figure 3B**). Thus, in addition to its well-characterized function in optimizing the peptide repertoire, our study demonstrates that tapasin alters the specificity of peptide-binding to HLA class I proteins near the C-terminus of HLA class I-bound peptides. For HLA-B*44:05, tapasin facilitates the inclusion of C-terminal tryptophan residues. Recent structural studies indicate the presence of a relatively open conformation of the MHC class I peptide binding groove in the tapasin-MHC class I complexes compared to the peptide-filled MHC class I complexes (11, 12). The F-pocket, formed by the α_2-1_ domain of the HLA class I peptide-binding groove, accommodates C-terminal residues of peptides (14, 34). The editing loop at the N-terminus of tapasin sits on top of the F pocket, while a β hairpin element of tapasin contacts the base of the F pocket, resulting in a “clamp” like binding of the tapasin to the α_2-_ι domain of the MHC class I peptide binding groove (11, 12). The editing loop of tapasin has been suggested to stabilize the widening of the F pocket creating a peptide-receptive conformation and tapasin also shields the C-terminus of an incoming peptide ligand (12). The increased detection of B*44:05 peptides with C-terminal tryptophans in the presence of tapasin suggests that tapasin-mediated widening of the F pocket is important to facilitate the binding of bulky amino acids such as tryptophan. Whether this is generally true for other HLA class I proteins that are permissive to tryptophans at the peptide-C-termini remains to be studied. The effects of tapasin on the peptidome composition and peptide-binding specificities of individual HLA class I proteins are undoubtedly linked both to the intrinsic specificity of peptide-binding to the particular HLA class I protein, as well as to the intrinsic flexibility and “openness” of the peptide-binding site in the peptide-free conformation. Tryptophan incorporation might only be facilitated by tapasin for those HLA class I allotypes that are both permissive for peptide C-terminal tryptophans and that have binding grooves that lack an intrinsic openness sufficient for tryptophan accommodation. Nonetheless, based on these studies, we predict that tapasin will generally influence the C-terminal amino acid compositions of HLA class I-bound peptides, although the specific influences will be HLA class I allotype-dependent.

We have observed significantly lower surface expression of B*44:02 compared to B*44:05 (**Figure 5A**) in some cell types even under tapasin-sufficient conditions (for example (31)). Despite the differences in surface expression, the surface stabilities of B*44:02 and B*44:05 were similar (**Figures 5B and 5C**), as reported previously (31). Lower SE values were measured for peptides identified for B*44:02 (**Figure 6C**) with an additional constraint at the P_C-2_ position, not seen for peptides identified for B*44:05 (**Figure 6C**). The presence of D116 in HLA-B*44:02 could favor the formation of salt bridges with positively charged peptide residues at the P_C-2_ position (**Figure 6E**). Despite the observed variations in amino acid preferences at the C-terminus of peptides (**Figure 6D**), the predicted binding affinities of B*44:02 peptides were similar among the unique and shared (with B*44:05) groups, indicating that the two groups are similarly optimized (**Figure 8B**). In contrast, the predicted affinities of B*44:05 peptides were significantly higher in the shared (with B*44:02) group compared to peptides unique to B*44:05 (**Figure 8A**). These findings suggest that B*44:05, in the presence of tapasin, acquires peptides via both the tapasin-dependent and tapasin-independent mechanisms. Peptides shared with B*44:02 have higher predicted affinities, consistent with a tapasin-dependent mode of loading. The presence of tapasin, thus partially optimizes peptide binding to tapasin-independent HLA class I allotypes, whereas optimization is applied more stringently to tapasin-dependent allotypes. Molecular dynamics simulations indicate that the peptide-binding groove of tapasin-independent allotypes exists in a partially closed conformation even in the peptide-free state, stabilized by the intrinsic structural characteristics of these proteins (35, 36). In contrast, the peptide-binding groove of tapasin-dependent allotypes exists in a more unstable and open conformation in the absence of a peptide ligand (35, 36). Tapasin-binding near the F-pocket of the peptide binding groove stabilizes peptide-receptive open conformations (12). A partially open conformation of tapasin-independent allotypes could facilitate the intrinsic binding of some peptides, but exclude the binding of other peptides, particularly those with bulky hydrophobic residues such as tryptophans, which could require tapasin’s assistance for loading, consistent with the data of **Figures 6 and 7**.

Overall, our study provides important insights into how tapasin shapes the peptidomes of both tapasin-dependent and tapasin-independent HLA class I allotypes, altering both the affinities and specificities of peptide binding. It is likely that additional low-abundance and low-affinity peptides are present under the tapasin-deficient conditions, which are undetectable using the current approaches. These studies also show why tapasin-independent allotypes are more protective in the context of infections (16). The abilities of tapasin-independent allotypes to load peptides under suboptimal assembly conditions could confer an important means of maintaining CD8^+^ T cell immunity in disease.

## Materials and methods

### Materials

PureProteome Protein A magnetic beads were obtained from EMD Millipore (LSKMAGA10). The RPMI-1640 media with L-glutamine, 1X PBS, L-glutamine, fetal bovine serum (FBS) and Antibiotic-Antimycotic (100X) were procured from Gibco (ThermoFisher Scientific). Several lab chemicals including brefeldin A, polybrene, phenylmethylsulfonyl fluoride (PMSF), sodium deoxycholate, octyl-beta-D glucopyranoside, iodoacetamide, triton X-100, triethanolamine (TEA), tris-base, glycine, protease inhibitor cocktail (catalog # P8340) and sodium azide were obtained from Sigma Aldrich. Dimethyl pimelimidate (DMP, catalog # 21667), FITC, acetonitrile, blasticidin S-HCl and trifluoroacetic acid were purchased from ThermoFisher Scientific. Bio-Rad 4–20% precast polyacrylamide gels (catalog # 4561096) were used for SDS-PAGE wherever manual gels were not used. Accugene 0.5 M EDTA solution was obtained from Lonza Bioscience.

### Cell lines and vector constructs

The 721.221 cell line (obtained from R. DeMars) lacks endogenous HLA-A, B, and C expression (37). The 721.221 cell lines expressing HLA-B*44:02 or HLA-B*44:05 (gifted by Dr. Wilfredo F. Garcia-Beltran, Ragon Institute) were grown in R10 media (RPMI 1640 (L-Glutamine containing media) supplemented with 10 % (v/v) FBS, 2 mM L-glutamine, and 1X antibiotic-antimycotic). The plasmids plentiCRISPRv2-BLAST, psPAX2 and pMD2.G were procured from Addgene.

### Crosslinking of Protein A magnetic beads to W6/32 antibody

The W6/32 antibody (20) was purified from mouse ascites and crosslinked to PureProteome Protein A magnetic beads as previously described (38). Briefly, following equilibration with 200 mM TEA pH 8.3, Protein A magnetic beads were incubated with the W6/32 antibody (2 μg antibody/μL of beads slurry) in 1X PBS for 1h at 4 °C. The unbound antibody was removed by washing the beads thrice with 200 mM TEA pH 8.3. Beads were crosslinked to the bound antibody using freshly prepared 5 mM DMP in 200 mM TEA pH 8.3 for 1h at ambient temperature. The crosslinking reaction was quenched using 5% (v/v) 1 M Tris-HCl pH 8.0 for 30 min at ambient temperature. Non-crosslinked antibody was further removed by incubating the beads with 0.1 M glycine-HCl pH 2.7 for 1 min. The beads were washed thrice with ice-cold PBS and stored at 4 °C in 1X PBS with 0.01% sodium azide.

### CRISPR/Cas9-based gene editing, single-cell cloning, and immunoblotting

To create 721.221-B*44:05 cells with tapasin knockout, the single guide RNA (sgRNA) targeting a sequence 5’-AAGCGGCTCATCTCGCAGTG-3’ within exon3 of the *tapasin* gene was used. The oligos for sgRNA synthesis (ordered from IDT) were cloned into the pLentiCRISPRv2-BLAST (pLB) vector. HEK293T cells were transfected with pLB-tapasin sgRNA vector together with the packaging plasmid psPAX2 and VSV-G envelope expressing plasmid pMD2.G. The virus particles were harvested, filtered, and used for infection of 721.221-B*44:05 cells by spinoculation at 2500 rpm for 2h at ambient temperature in the presence of 8 μg/mL polybrene. Following the selection of transduced cells with 5 μg/mL blasticidin, single-cell cloning was performed by plating the cells at a density of 5 cells/mL in 96 well plates.

For detection of tapasin knockout, single-cell clones were harvested and lysed in 1X tris buffered Saline (TBS) with 1% triton X-100. Tapasin was detected in the lysates following 12% SDS-PAGE and western transfer using anti-tapasin antibody (MABF249 from Sigma-Aldrich).

HLA-B heavy chain and β2-microglobulin (β2m) were probed using HC10 (mouse hybridoma procured from University of Michigan Hybridoma Core) and purified anti-β2m (Biolegend; catalog # 316302) antibodies.

### Flow cytometry staining and antibody binding capacity determination

Before each immunoaffinity purification experiment discussed in the following sections, the cells were harvested and stained for surface HLA class I using W6/32-FITC antibody (FITC conjugation was performed in the lab following the manufacturer’s protocol). The staining was done in 1X PBS containing 2% FBS (staining buffer) for 30 min at 4 °C. The cells were washed twice with the staining buffer and stained with 7-AAD viability dye. Unfixed cells were analyzed by flow cytometry using BD LSRFortessa™ cell analyzer.

For quantitative flow cytometry experiments Quantum™ simply cellular anti-mouse IgG beads (Bangs Laboratory, Inc.) were stained concurrently with the cells using the same antibody dilution and analyzed at the same time as the cells.

The analysis of flow cytometry data was performed using FlowJo™. Live cells were gated based on negative 7-AAD staining and W6/32 geometric mean fluorescence intensity (gMFI) values were determined. Standard curves were plotted using the geometric mean fluorescence intensity (gMFI) values for various bead populations *vs*. the known number of Fc receptors on the respective beads as given by the manufacturer. The antibody binding capacity (ABC) of cells was determined by interpolation from standard curves. The W6/32-ABC values of cells represent the approximate number of HLA class I molecules expressed on individual cells.

### Stability and half-life calculations of cell surface peptide-HLA-B complexes

Cell surface stability of peptide-HLA-B complexes was measured as described previously (24, 39). Briefly, the 721.221-B*44:05 cells (wild-type or tapasin-KO) and 721.221-B*44:02 cells were counted and added to a 96-well plate in triplicates for each time point in RPMI+10% FBS media. Brefeldin A was added at 0.5 μg/mL concentration and the cells were harvested after incubation in brefeldin A for indicated time periods. The cells were washed with 1X PBS and incubated with W6/32-FITC antibody for 30 minutes at 4 °C in the staining buffer mentioned above followed by two washes in the staining buffer. Cells were stained with viability dye 7-AAD and analyzed by flow cytometry using BD LSRFortessa™ cell analyzer.

The geometric MFI values for cells at various time points were analyzed in GraphPad Prism using one phase decay model constraining the plateau to a constant value of zero to determine the half-life values. Average half-lives from multiple experiments were analyzed by two-tailed unpaired t-tests to determine significance values.

### Immunoaffinity (IA) purification of peptide-HLA class I complexes

For immunoaffinity purification, as published previously (21), 721.221-B*44:02 and 721.221-B*44:05 (wild type and tapasin-KO cells) were grown to ~2 to 4 x 10^8^ cell density. The cells were lysed in 10 mL of lysis buffer (1 mM EDTA, 1 mM PMSF, 1% octyl-beta-D glucopyranoside and 0.25 % sodium deoxycholate in 1X PBS supplemented with protease inhibitor cocktail (1:200) and freshly prepared 0.2 mM iodoacetamide). The lysis was performed at 4 °C for 1 h on rotation. The lysates were cleared by centrifugation at 14000 rpm for 30 min at 4 °C. The cleared lysates were incubated with 2 mL blank protein A beads for pre-clearing at 4 °C for 30 min on rotation. The pre-cleared lysates were incubated with 2 mL of protein A beads crosslinked to W6/32 antibody (W6/32 beads) on rotation for 3h at 4 °C. The beads were harvested and washed with 8 column volumes of 1X PBS followed by 1 column volume each of 0.1X PBS and water. HLA class I complexes were eluted in six serial fractions with 0.2 mL of freshly prepared 0.2% (v/v) trifluoracetic acid. For reuse, the beads were regenerated with 1 column volume of water and 3 column volumes of 1X PBS before storing the beads in 1X PBS with 0.01% sodium azide. All the wash buffers as well as the elution buffer were kept ice-cold.

### Isolation of peptides

The peptides were further isolated from the immunoprecipitated peptide-HLA class I complexes using either Sep-Pak C18 cartridges (Waters Corp, Catalog # WAT020515) or stage tips packed manually using C18 extraction disks (20 stacked disks, 3M™ Empore™). The stage tips or the cartridges were activated with 80% acetonitrile in 0.1% TFA followed by a wash with 0.1% TFA. The eluates were loaded on the stage tips or Sep-Pak cartridges followed by washing with 0.1% TFA. Peptides were eluted using 30% acetonitrile in 0.1% TFA and dried using vacuum centrifugation.

### Mass spectrometry

The peptides were resuspended in 2% ACN, 0.1% formic acid for mass spectrometry. The mass spectrometry was performed in Proteomics Resource Facility in the department of pathology at the University of Michigan on Thermo Q Exactive HF. For MS/MS, 2 μL of peptide solution was resolved on a nano-capillary reverse phase column (Acclaim PepMap C18, 2 microns, 50 cm, ThermoScientific) using 0.1% formic acid/acetonitrile gradient at 300 nL/min (2-25% acetonitrile for 105 min; 25-40% acetonitrile for 20 min followed by a 90% acetonitrile wash for 10 min and a further 30 min re-equilibration with 2% acetonitrile) and directly introduced into *Q Exactive HF* mass spectrometer (Thermo Scientific, San Jose CA). MS1 scans were acquired at 60K resolution (AGC target=3e^6^, max injection time=50ms). Data-dependent high-energy C-trap dissociation MS/MS spectra were acquired for the 20 most abundant ions (Top20) following each MS1 scan (15K resolution; AGC target=1e^5^; relative collision energy ~28%).

### Mass spectrometry data analysis

All MS raw files (.raw files) were analyzed using PEAKS studio software. MS/MS spectra were searched against a sequence database created from the human sequences of UniprotKB/Swiss-Prot (Download date: 2015-06-19) appended with reversed protein sequences as decoys and common contaminants. The mass tolerance was set at 10 ppm for precursor ions and 0.02 Da for the fragment ions. Isotopic error correction and common variable modifications of methionine oxidation, N-terminal acetylation, and cysteine carbamidomethylation were enabled. The enzyme specificity was set as ‘none’. The false discovery rate (FDR) of 1% was used filtration of peptide-spectra matches (PSMs). MS/MS data from six runs of peptides eluted each from the B*44:02, B*44:05 -WT, and B*44:05-tapasin-KO cells were used for analyses. The peptides of lengths 8-14 were included in further analysis and the selected peptides were further filtered to remove 1) common contaminant peptides, 2) peptides derived from bovine proteins, and 3) peptides present in the MS/MS run of HLA class I null 721.221 cells.

### Analyses of peptidome datasets

The mass spectrometry outputs provided by the PEAKS studio software were analyzed using an R language-based script that converted individual cells with numerical outputs into “TRUE” and “FALSE” labels, such that any peptide that was detected in a mass spectrometry run appeared as “TRUE”, whereas undetected peptides appeared as “FALSE.”

The peptides were then grouped based on the number of runs across which each peptide was detected

Grouping of the peptides involved dividing peptides into shared and unique groups. Peptides unique to a specific allotype or condition were those detected in ≥2 independent MS/MS runs only for that allotype or condition. Peptides shared between any two conditions were ones detected in ≥2 independent MS/MS runs of each of the two conditions under consideration.

### Shannon entropy plots, Seq2Logo plots, and prediction of binding affinities of peptides

Shannon Entropy (SE) was used as a diversity index to determine the amino acid variabilities at specific positions within a peptide sequence for each allele or condition (23, 24). The peptide sequences from each dataset were assessed using equation “***E***(***i***) = -***∑qi***(***log***_**2**_***qi***) here *E(i*) is the SE for position *‘i’* within the peptide sequence and *qi* is the probability of finding an amino acid at position *‘i’* The maximum SE = 4.3 indicates maximum flexibility when all the 20 amino acids are found at a position with equal probability. The smaller the value of SE for a position, the higher the restrictiveness of that position. Individual SE values were calculated for each position within a subset of shared or unique peptides grouped by length.

The consensus Seq2Logo motifs (26) for individual peptide subsets, for example, shared and unique peptides for each allele or condition were plotted using a web-based sequence logo generation tool (https://services.healthtech.dtu.dk/service.php?Seq2Logo-2.0). The binding affinities of peptides within the shared and unique subsets were predicted using the NetMHCpan-4.1 (27) server (https://services.healthtech.dtu.dk/service.php?NetMHCpan-4.1).

### Amino Acid Distribution Calculations

The fractional distribution of amino acids at the P_C_ and P_C-2_ positions for the 9-mer, 10-mer, and 11-mer peptides was calculated from the fraction of total peptides in a subset with a particular amino acid at a specific position. These fractional distributions were plotted using Prism GraphPad Prism 9.

### Statistical analysis

Statistical analyses were performed in GraphPad Prism version 9.4.1 or using R studio via the specific tests/methods mentioned in the respective figure legends.

## Acknowledgements and funding sources

This work was supported by the National Institutes of Health Grants to MR (R01AI044115 and R21AI164025), AJZ (T32AI007528 and T32AI007413) and AIN (R01GM94231), and National Cancer Institute grant to AIN (U24CA271037). We thank Elizabeth Smith from the University of Michigan Hybridoma Core for antibody production. We thank Corey Powell from CSCAR (Consulting for Statistics, Computing and Analytics Research) at University of Michigan for help with the statistical analysis.

## Author Contributions

Conception and Design, MR, AJZ; Acquisition of Data, AJZ, AK, VB; Analysis and Interpretation of Data, AS, MBM, AJZ, AK, VB, AIN, MNC, IG, MR; Writing, MR, AS, AK, MBM, AJZ, VB; Editing, AS, AK, AJZ, VB, AIN, MNC, MR; Statistical analysis, AS, AK, MBM, MC, MR; Funding Acquisition, MR, AIN; Supervision, MR

## Conflicts of Interest

All the authors have no conflicts of interest to declare.

## Notes

### Competing Interest Statement

The authors have declared no competing interest.

