## Supplementary files for "Mass spectrometric profiling of HLA-B44 peptidomes provides evidence for tapasin-mediated tryptophan editing"

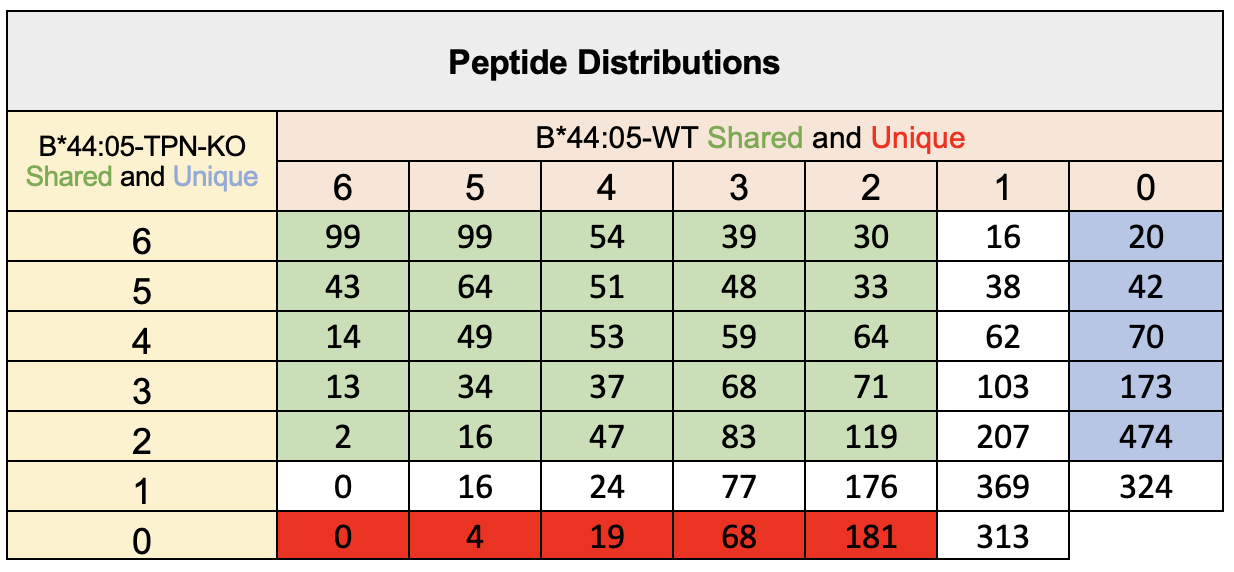

**Supplementary table 1:** Table shows the distribution of the full set of peptides identified from six MS/MS runs grouped into those unique to B*44:05-WT (red), those unique to B*44:05-TPN-KO (blue) and those shared across both B*44:05-WT and B*44:05-TPN-KO conditions (green).

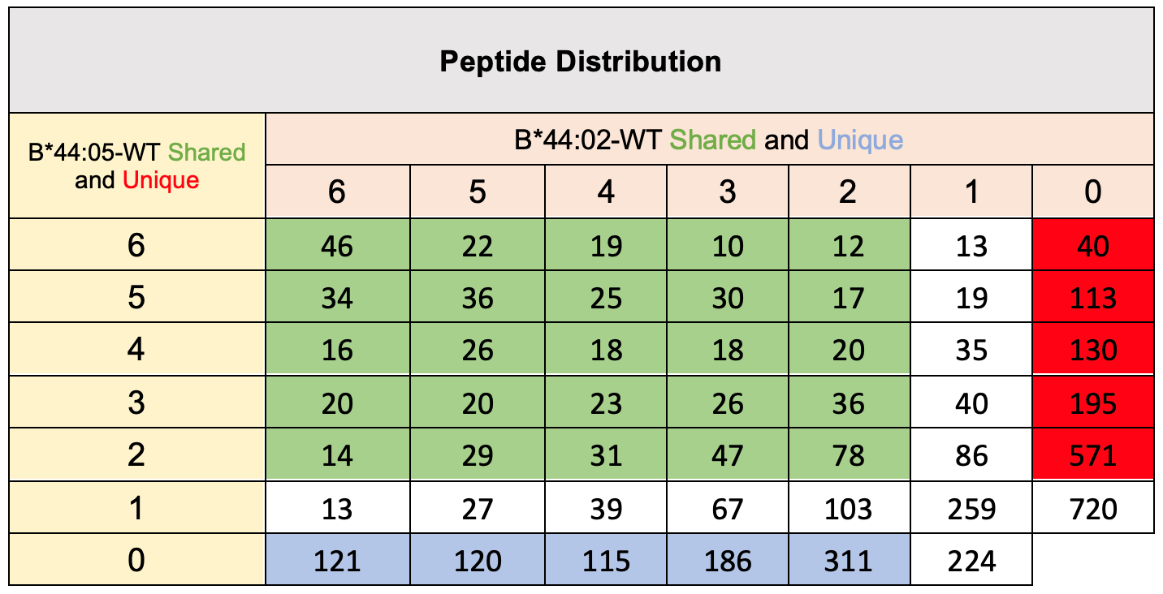

**Supplementary table 2:** Table shows the distribution of the full set of peptides identified from six MS/MS runs grouped into those unique to B*44:05-WT (red), those unique to B*44:02-WT (blue) and those shared across both B*44:05-WT and B*44:02-WT conditions (green).
